# Diversification of biofilm architecture among freshwater *Pararheinheimera* isolates

**DOI:** 10.64898/2026.02.25.707958

**Authors:** Amanda E. Appel, Alexandra G. Goetsch, Ellen W. van Wijngaarden, Daniel J. Novacek, Peter M. Burnham, Meredith N. Silberstein, David M. Hershey

## Abstract

Air-liquid interfaces (ALIs) at the upper layer of oceans, lakes and rivers cover the majority of the earth’s surface. Microbes are known to accumulate at these resource-rich boundaries, but the mechanisms of ALI colonization are often assumed to mirror the formation of pellicle biofilms by non-aquatic organisms. Here, we analyzed ALI colonization by natural aquatic bacteria. We used samples from a freshwater lake to enrich for microbes that colonize the ALI in liquid growth medium. Mixed-species pellicles formed rapidly in these enrichments, were structurally stable for weeks and displayed a pronounced ecological succession. We isolated 31 members of the genus *Pararheinheimera* from early stages of mixed-species pellicle maturation. Five phylogenetically distinct *Pararheinheimera* clades were identified, each with a shared colony morphology. We used representative isolates to show that only one *Pararheinheimera* clade formed thin, adherent films at the ALI resembling classical pellicles. Isolates from the four remaining clades formed floating structures that could be categorized either as non-adhesive films or large viscous masses. Viscous mass (VM) pellicle formation was a polyphyletic trait that correlated with a highly mucoid appearance on agar plates, suggesting that the process is driven by copious secretion of extracellular matrix. Matrices from VM biofilms were largely non-adhesive, contained a mixture of acidic polysaccharides and proteins and formed thermally stable, shear-thinning hydrogels. Our results demonstrate that ALI colonization strategies vary widely even among closely related aquatic bacteria and identify VM pellicles as a distinct biofilm architecture with unique mechanical properties.

**Importance:** Lakes, rivers and oceans contain a boundary between the air and the water’s surface known as the air-liquid interface (ALI). Microbial communities that populate the ALI play crucial roles in nutrient cycling, but how aquatic microbes partition to these sites remains poorly characterized. Our study investigated how bacteria from a freshwater lake accumulate at the ALI. Lake water samples incubated in nutrient medium formed a layer of cells known as a pellicle biofilm at the ALI, and we isolated 31 different bacteria from a genus (*Pararheinheimera*) that was abundant during the early stages of pellicle formation. Only a subset of *Pararheinheimera* isolates formed traditional pellicle biofilms. Most formed either thin, non-adhesive films or large, gelatinous aggregates that appeared to persist at the ALI due to buoyancy. These findings expand our understanding of biofilm diversity in aquatic systems and suggest that the production of buoyant hydrogels may play an important role in structuring microbial communities at air–water boundaries.

## Introduction

Microbes form assemblies known as biofilms that contain living cells embedded in an extracellular matrix(1). The biofilm lifestyle is more prevalent in most ecosystems than the free-living, planktonic state traditionally associated with bacteria(2). Cells in biofilms benefit from enhanced access to nutrients, protection from environmental stressors, and opportunities for social interactions. Biofilms also play central roles in processes ranging from nutrient cycling and carbon sequestration(3) to plant-microbe interactions(4), chronic infections(5) and industrial biofouling(6). These activities drive the broad ecological relevance of biofilms and their profound impact on society(7).

Extracellular matrices serve a crucial role in biofilm function. Secreted polysaccharides, proteins and extracellular DNA combine with water, ions and adsorbed organic compounds from the environment to create a complex scaffold(8). This matrix displays both solid- and liquid-like features, resulting in an emergent property known as viscoelasticity. Viscoelastic behavior enhances surface adhesion, limits permeability to exogenous compounds and provides resilience to mechanical forces such as shear flow(9). Biofilm matrix composition varies widely among bacteria, particularly in the chemical structures of secreted exopolysaccharides (EPSs)(10, 11). The resulting differences in elasticity, cohesiveness, porosity and other properties enable biofilms with diverse architectures to thrive across a wide range of ecological niches(12).

The classical model of biofilm formation describes a stepwise program in which planktonic cells attach to a solid surface, secrete a biofilm matrix as they proliferate and eventually disperse into bulk liquid(13). Much of this framework emerged from pioneering studies with model organisms such as *Pseudomonas aeruginosa, Vibrio cholerae, Bacillus subtilis* and *Escherichia coli*(14–17). These traditional models have yielded deep insights into the signaling networks that control biofilm development(18), the composition of the extracellular matrix(19) and strategies for treating biofilm-associated infections(20). However, the classic model is rooted in attachment to solid surfaces. It does not fully capture the diversity of biofilm architectures observed in natural environments(21).

Aquatic ecosystems contain a distinctive niche at the air-liquid interface (ALI), where nutrients, oxygen, light, and other resources are readily available(22–24). Many bacteria can form biofilms known as pellicles at the ALI when incubated in liquid growth medium without agitation(25). Bacterial strains that display rugose or wrinkled colony morphologies on agar plates tend to form structures called physically cohesive (PC) pellicles (26). These structures behave as mechanically continuous films coating the ALI and often display high structural stiffness(26–28). Matrix composition and genetic pathways controlling PC-type pellicle formation largely overlap with those of traditional solid-surface biofilms(29–32), suggesting that this architecture reflects an extension of the surface-associated biofilm program rather than a distinct, surface-independent strategy.

Physically cohesive (PC) pellicle formation has been observed in certain soil isolates, human pathogens and evolved or engineered mutants with enhanced biofilm formation(33, 34, 29, 35, 36, 27, 37, 38). ALI colonization is not thought to represent an ecologically relevant behavior for many of these organisms. Despite extensive evidence that microbes at the ALI play crucial roles in aquatic ecosystems, pellicle formation in natural aquatic isolates has rarely been investigated in detail (39, 31). Fundamental knowledge of how microbes access this niche is lacking. The biofilm matrices of aquatic bacteria likely display unique properties that allow cells to balance buoyancy, surface tension, and shear flow in order to persist at the surface of the water column. Examining pellicle biofilms specifically in aquatic isolates has the potential to illuminate these mechanisms.

Here, we describe a group of freshwater bacteria that forms unusual biofilms at the ALI. Water from a eutrophic lake in Madison, WI was used to enrich mixed-species pellicle biofilms in liquid growth medium. 31 different strains from the genus *Pararheinheimera* were isolated during the early stages of pellicle formation in these enrichment cultures, and we grouped these isolates into five distinct phylogenetic clades. Individual *Pararheinheimera* isolates formed biofilms at the ALI with a range of architectures, including physically cohesive films, non-adhesive layers and large viscous masses. Extracted matrices from the floating, viscous mass (VM)-type biofilms contain an acidic exopolysaccharide and form thermostable, shear-thinning hydrogels in aqueous solution. These results highlight the diverse biofilm architectures present among aquatic microbes and suggest that buoyancy may play a key role in structuring biofilms at natural ALIs.

## Results

### Mixed-species pellicles enriched from lake water display an ecological succession

We evaluated the ability of freshwater microbes to colonize the air–liquid interface (ALI) by inoculating water samples from Lake Mendota (Madison, WI) into tubes containing liquid medium. These cultures developed a visible layer of biomass at the ALI when incubated statically (Figure 1A). Biofilms appeared within two days and remained stable for 15–20 days, after which they sank to the bottom of the tubes. Biomass was collected from the ALI at defined timepoints using the wide end of a pipette tip. ALI samples displayed adhesive properties when transferred to glass slides and stained with crystal violet (Figure 1B). Biomass transferred one day after inoculation produced faint and uneven staining. Transferred biomass matured into a uniform and strongly adherent layer on days 2, 4, and 8. In contrast, biomass collected at day 16 showed reduced adhesion and a patchy staining pattern consistent with decreased cohesion. Phase-contrast microscopy of ALI material transferred to agarose pads (Figure 1C) showed that microcolonies formed within 24 hours and matured into a dense cellular matrix by day 4. This dense structure persisted through the remainder of the time course. Cells within these microcolonies were predominantly rod-shaped or coccoid, with substantial variation in length and aspect ratio. Single cells were difficult to resolve in samples collected at days 4, 8, and 16 due to the thickness of the biofilms.

**Figure 1:**
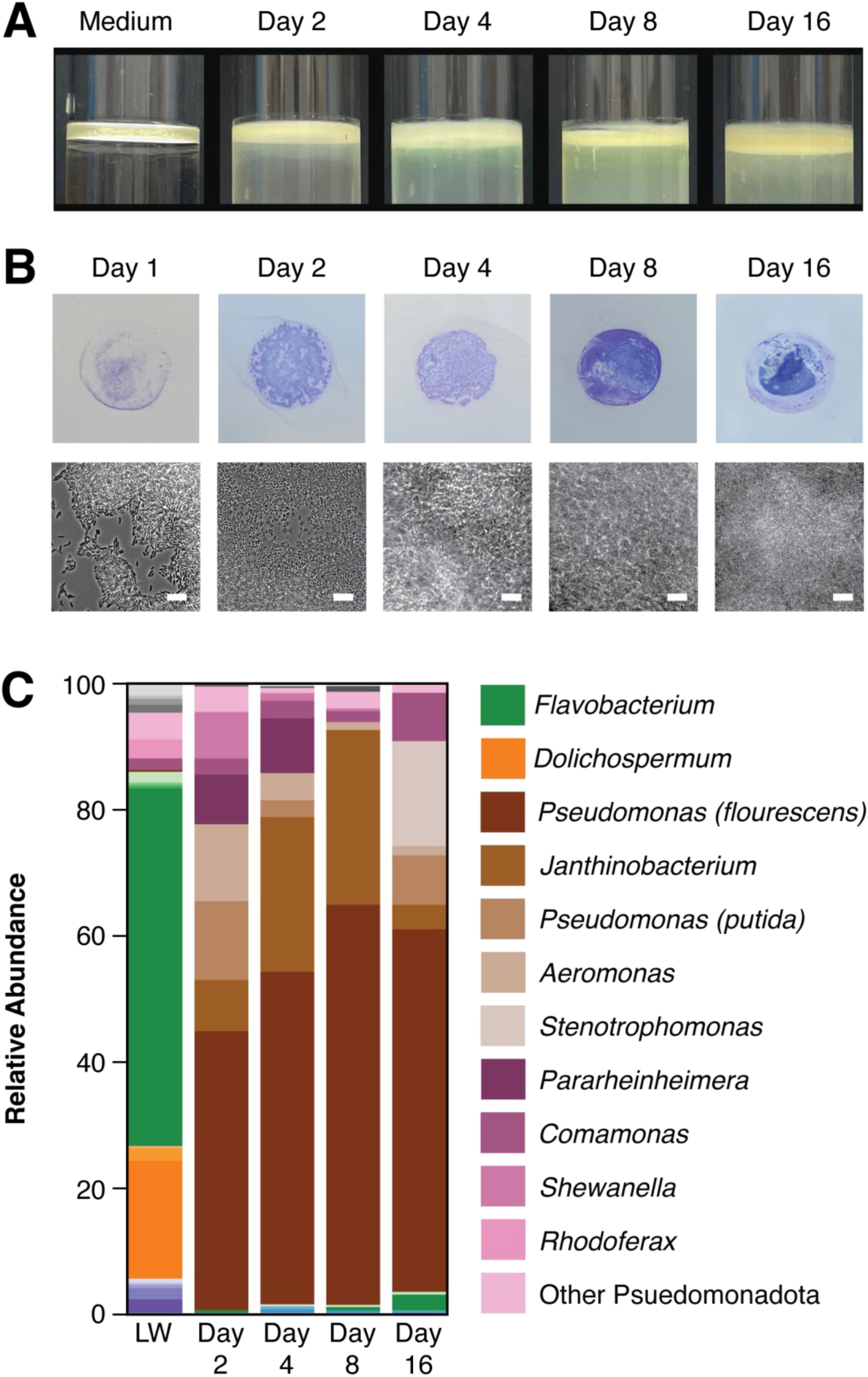
Enrichment of mixed-species pellicles from natural lake water. (A) Pellicles form at the ALI when lake water is incubated statically in PYE medium. A layer of biomass can be seen at the ALI within two days and persists throughout a 16-day incubation. (B) Extracted ALIs form adhesive mats. Biomass from the ALI was extracted with a pipette tip and transferred to glass for crystal violet staining (top) or agarose pads for microscopy (bottom). Adhesive biomass was observed after 24 hours and a homogenous adhesive layer was present by 2 days. Microcolonies formed within 24 hours and merged into a single layer by 4 days. Scale bars represent 1.25µm. (C) 16S amplicon sequencing analysis of bacteria present during mixed-species pellicle formation shows an ecological succession. *Pseudomonada* dominate enriched pellicles throughout a 16 day time-course. *Pseudomonas* strains make up more than half of reads at all time points. Early (*Shewanella, Pararheinheimera*), middle (*Aeromonas*, *Janthinobacterium*) and late (*Stenotrophomonas*) accumulating genera can be identified among the remaining reads. Clades that compose greater than 5% of reads in at least one sample are highlighted in the key.

We used 16S rRNA gene amplicon sequencing to monitor changes in community composition during ALI colonization (Figure 1D). Initial lake water samples were dominated by Flavobacteria and cyanobacteria from the genus *Dolichospermum*. Proteobacteria from genera such as *Comamonas* and *Rhodoferax* were present at lower but detectable levels. Proteobacteria dominated the ALI after inoculation into liquid medium. By two days post inoculation, Proteobacteria accounted for more than 98% of reads, while Flavobacteria and *Dolichospermum* were no longer detected. Proteobacteria remained above 95% of the ALI community throughout the 16-day time course.

The Proteobacterial population at the ALI underwent pronounced shifts during biofilm development. *Pseudomonas* strains comprised more than half of all reads at the ALI throughout the experiment despite representing less than 0.5% of sequences in the initial lake water inoculum. *Pararheinheimera* and *Shewanella* strains each bloomed to nearly 10% of the community at day 2 before declining to undetectable levels by day 8. *Aeromonas* exhibited a similar early increase but stabilized at approximately 1% of the population at days 8 and 16. *Janthinobacterium* peaked at 28% of reads at day 8 before declining thereafter. In contrast, *Stenotrophomonas* strains accumulated only at late stages of biofilm development, comprising 17% of reads at day 16 but remaining undetected at earlier timepoints. Together, these results show that freshwater microbial communities rapidly assemble dynamic and compositionally structured biofilms at the air–liquid interface when exposed to nutrient-replete conditions.

### Pararheinheimera isolates from mixed-species pellicles form five sub-clades

We subjected water samples from Lake Mendota to the enrichment procedure described above under different nutrient conditions. Untreated lake water and lake water supplemented with various concentrations peptone-yeast extract (PYE) medium were incubated for 16 days. Samples from the ALI were collected throughout each enrichment and serially diluted on solid medium to obtain single colonies. Colonies representing a range of morphologies were isolated, and each was characterized by sequencing the 16S rDNA gene. Across all nutrient regimes, bacteria annotated as *Pararheinheimera* were consistently recovered from unamended lake water and from the earliest stages of ALI enrichment. Over a period of three years, we obtained 31 *Pararheinheimera* isolates from Lake Mendota (Table 1).

**Table 1:** *Pararheinheimera* strains isolated from mixed-species pellicle enrichments. Isolates were obtained over a four-year period from pellicles grown for 0-4 days in different nutrient regimes (see Materials and Methods). Though isolations from pellicle enrichments were continued for up to one month, *Pararheinheimera* was only recovered within the first four days of inoculation. The “RC#” column contains number from our internal lake isolate collection and are listed to facilitate strain requests.

| Isolate | RC# | Clade | Condition | Incubation time (days) | GenBank Accession |
| --- | --- | --- | --- | --- | --- |
| PP2D | 1 | 1 | PYE | 2 | PX310769 |
| PP2H | 2 | 1 | PYE | 2 | PX310770 |
| PP2J | 3 | 1 | PYE | 2 | PX310771 |
| PP4B | 4 | 1 | PYE | 4 | PX310772 |
| PP4D | 5 | 1 | PYE | 4 | PX310773 |
| PP4F | 6 | 1 | PYE | 4 | PX310774 |
| PP4I | 7 | 1 | PYE | 4 | PX310775 |
| B1H9 | 8 | 5 | 5% PYE | 1 | PX310747 |
| B1H13 | 9 | 5 | 5% PYE | 1 | PX310748 |
| B2H2 | 10 | 2 | 5% PYE | 2 | PX310749 |
| B2H7 | 11 | 5 | 5% PYE | 2 | PX310750 |
| B1H7 | 12 | 4 | 5% PYE | 1 | PX310746 |
| B2H12 | 13 | 1 | 5% PYE | 2 | PX310752 |
| DN2 | 14 | 3 | 5% PYE | 2 | PX310755 |
| DN4 | 15 | 3 | 0.5% PYE | 2 | PX310757 |
| C3L8 | 16 | 1 | 0.5% PYE | 3 | PX310753 |
| T2C2 | 17 | 1 | 0.5% PYE | 2 | PX310776 |
| C3L9 | 18 | 2 | 0.5% PYE | 3 | PX310754 |
| DN3 | 19 | 5 | 5% PYE | 2 | PX310756 |
| DN5 | 20 | 2 | 5% PYE | 4 | PX310758 |
| DN7 | 21 | 2 | Lakewater | 4 | PX310759 |
| DN8 | 22 | 2 | Lakewater | 4 | PX310760 |
| DN9 | 23 | 2 | Lakewater | 4 | PX310761 |
| DN11 | 24 | 2 | Lakewater | 4 | PX310763 |
| DN10 | 25 | 2 | Lakewater | 4 | PX310762 |
| DN17 | 26 | 3 | Direct plating | 0 | PX310764 |
| DN18 | 27 | 5 | Direct plating | 0 | PX310765 |
| DN19 | 28 | 3 | Direct plating | 0 | PX310766 |
| DN20 | 29 | 3 | Direct plating | 0 | PX310767 |
| DN21 | 30 | 3 | Direct plating | 0 | PX310768 |
| B2H8 | 31 | 4 | 5% PYE | 2 | PX310751 |

We aligned sequences of the 16S rDNA gene from all 31 *Pararheinheimera* isolates along with those from a previously described group of strains from the genera *Pararheinheimera, Rheinheimera, Alishewanella* and *Arsukibacterium*(40). A maximum-likelihood phylogenetic tree from this alignment largely supported the previously described distinctions among these four genera and confirmed the placement of all 31 Lake Mendota isolates within the *Pararheinheimera* genus (Figure 2A). The addition of dozens of new *Pararheinheimera* sequences allowed us to delineate five sub-clades. Four of these clades include representatives from the Sisinthy et al. analysis(40), but one group (clade II) includes only isolates from our collection.

**Figure 2:**
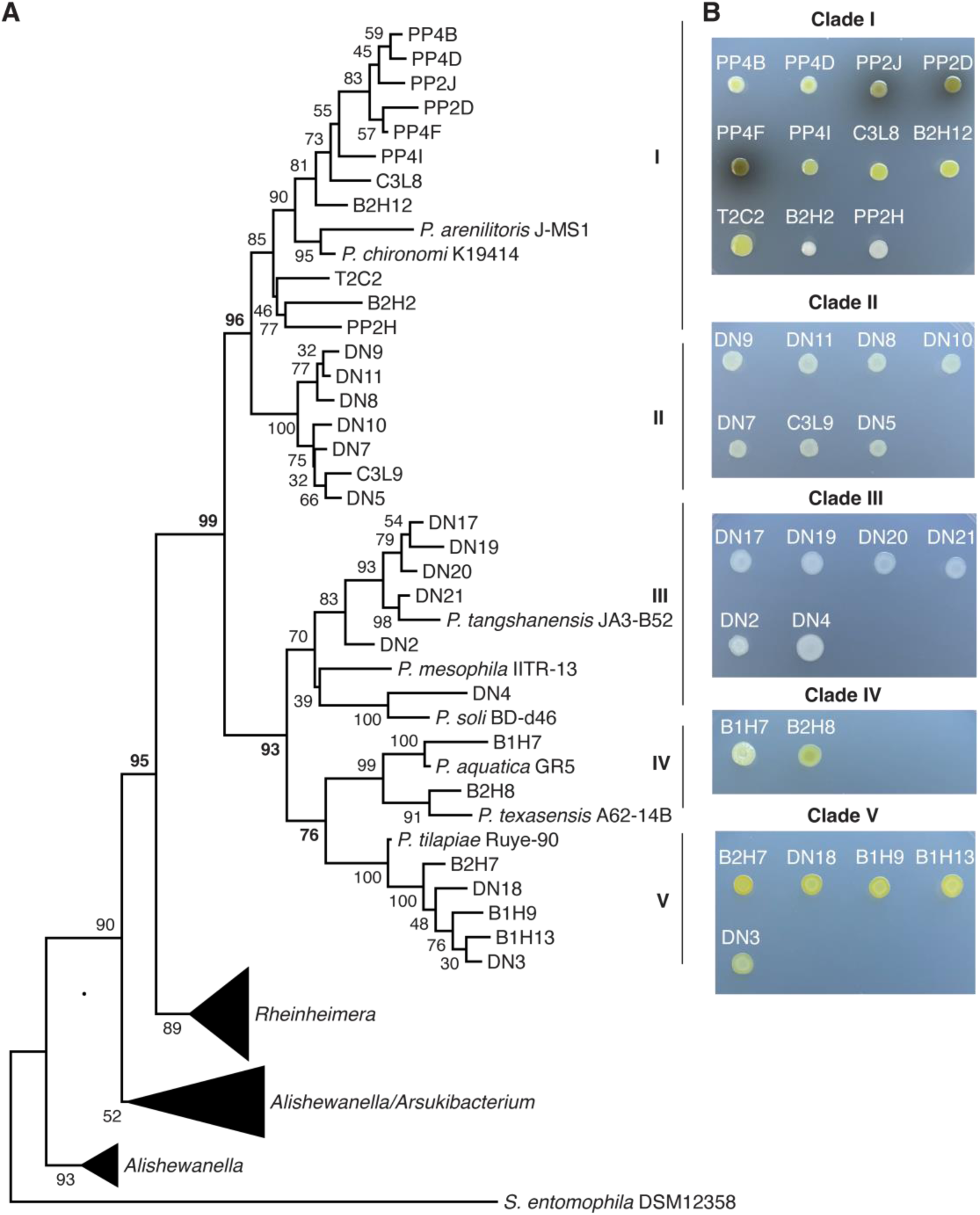
Freshwater *Pararheinheimera* isolates encompass five distinct sub-clades. (A) Maximum likelihood phylogenetic tree of 16S rDNA sequences obtained from *Pararheinheimera* isolates. Strains analyzed by Sisinthy et al. are included in the analysis for reference. Five sub-clades are identified. (B) Colony morphologies of *Pararheinheimera* on agar plates. Extensive similarities in colony appearance are observed among members of each sub-clade.

The assignment of these subclades is supported by visual analysis of colony morphology on agar plates (Figure 2B). Clade I isolates form yellowish colonies with highly mucoid appearance, and some clade I strains secrete a brown pigment. Clade II colonies appear white with slight wrinkling. Clade III isolates form white, non-mucoid colonies. The two representatives from clade IV have a faint yellow color. Both appear mucoid, though B1H7 colonies are more mucoid than those of B2H8. Clade V isolates form non-mucoid orange colonies. Morphological differences observed during growth on agar plates indicate that the five *Pararheinheimera* clades likely differ in their biofilm formation behavior despite their close phylogenetic relationships.

### Distinct biofilm architectures among Pararheinheimera subclades

One representative from each of the five *Pararheinheimera* clades was chosen for detailed analysis of ALI colonization (Figure 3). Pellicle formation in our five *Pararheinheimera* representatives was compared to three established ALI colonizers: *Caulobacter crescentus* CB15(31), *Pseudomonas protegens* Pf-5(41), and *Shewanella oneidensis* MR-1(42). Each strain was incubated statically in liquid medium to allow for pellicle development (Figure 3A). *Pararheinheimera sp.* B2H12 (clade I) and B1H7 (clade IV) formed a large gelatinous mass at the ALI. Strains DN11 (clade II) and DN20 (clade III) displayed cloudy menisci, suggesting the presence of cells at the ALI. Similar cloudy menisci were observed for CB15, Pf-5 and MR-1. Clade V representative B2H7 formed a thin film at the ALI that was highly pronounced due to the characteristic pigmentation of this strain (Figure 2). The culture tubes were washed and stained with crystal violet (CV) to determine whether each strain adhered to the glass surface (Figure 3B). Tubes from CB15, Pf-5 and MR-1 cultures stained in a ring at the site of the ALI, consistent with the formation of an adhesive pellicle layer. B2H7 (clade V) was the only *Pararheinheimera* representative that stained strongly. The clade IV representative B1H7 stained weakly, and representatives from clades I, II and III did not stain at all (Figure 3B).

**Figure 3:**
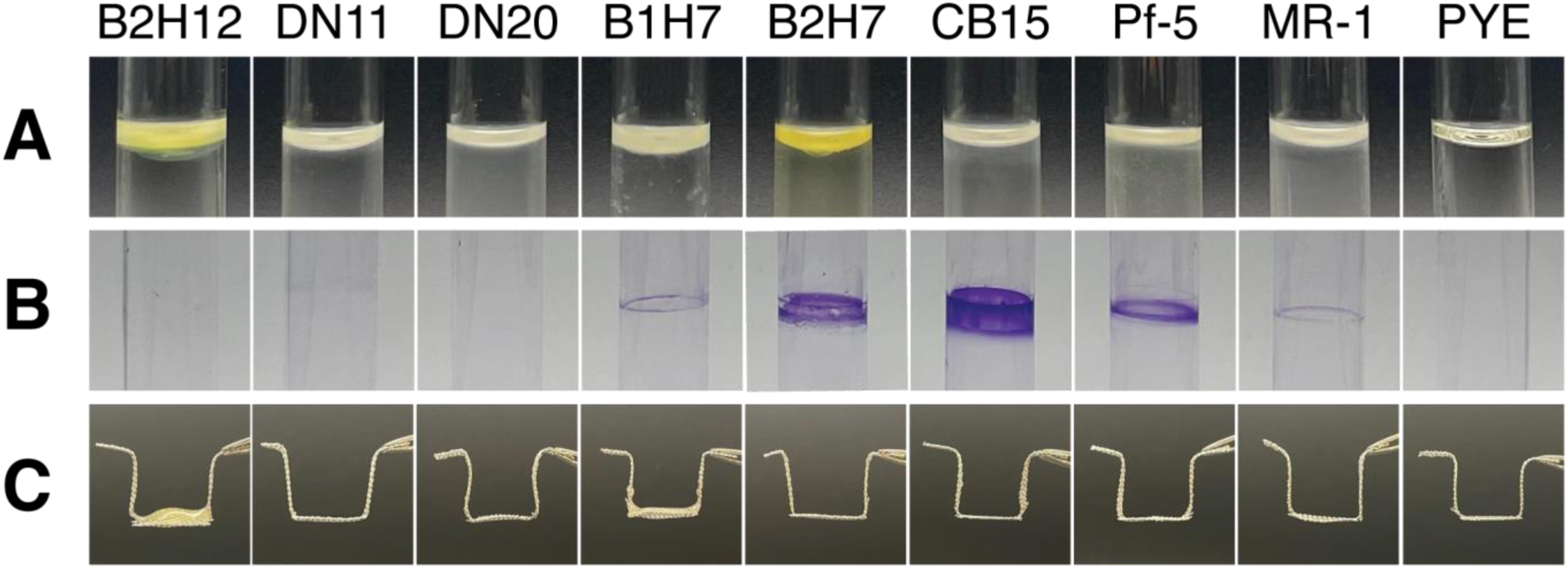
ALI colonization by *Pararheinheimera* isolates. (A) Images of ALIs from static liquid cultures of representative *Pararheinheimera* strains, along with known ALI colonizers *Caulobacter crescentus* CB15, *Pseudomonas protegens* Pf-5 and *Shewanella oneidensis* MR-1. (B) Crystal violet staining of static liquid cultures. Culture tubes were washed with water before staining to remove unattached biomass. (C) Biomass filtered with steel mesh shows the accumulation of gelatinous biomass in B2H12 (clade I) and B1H7 (clade IV) cultures.

Despite the lack of CV staining for B2H12 (clade I), DN11 (clade II) and DN20 (clade III), there were still visual indications of biomass at the ALI in these cultures. We modified our growth conditions to examine the possibility that biofilms accumulate at the surface without adhering to the culture vessel. Cultures were grown statically in 12-well plates with a layer of steel mesh suspended a few millimeters above the bottom of each well. Gelatinous biofilms were filtered from the ALI of B2H12 (clade I) and B1H7 (clade IV) cultures when the mesh was lifted (Figure 3C). Steel mesh filtrates from DN11 (clade II), DN20 (clade III), B2H7 (clade V) and all three established ALI colonizers (CB15, Pf-5 and MR-1) lacked obvious biomass. Finally, we directly visualized cells at the ALI in DN11 and DN20 cultures by using a pipette to transfer the ALI to an agarose pad for microscopy (Figure 4). Both cultures contained continuous films that resembled the high cell density films from B2H7, CB15, Pf-5 and MR-1. We conclude that representatives from all five *Pararheinheimera* clades accumulate at the ALI in static culture, though the resulting biofilm architectures (non-adhesive films, physically-cohesive films, large viscous masses) are distinct.

**Figure 4.**
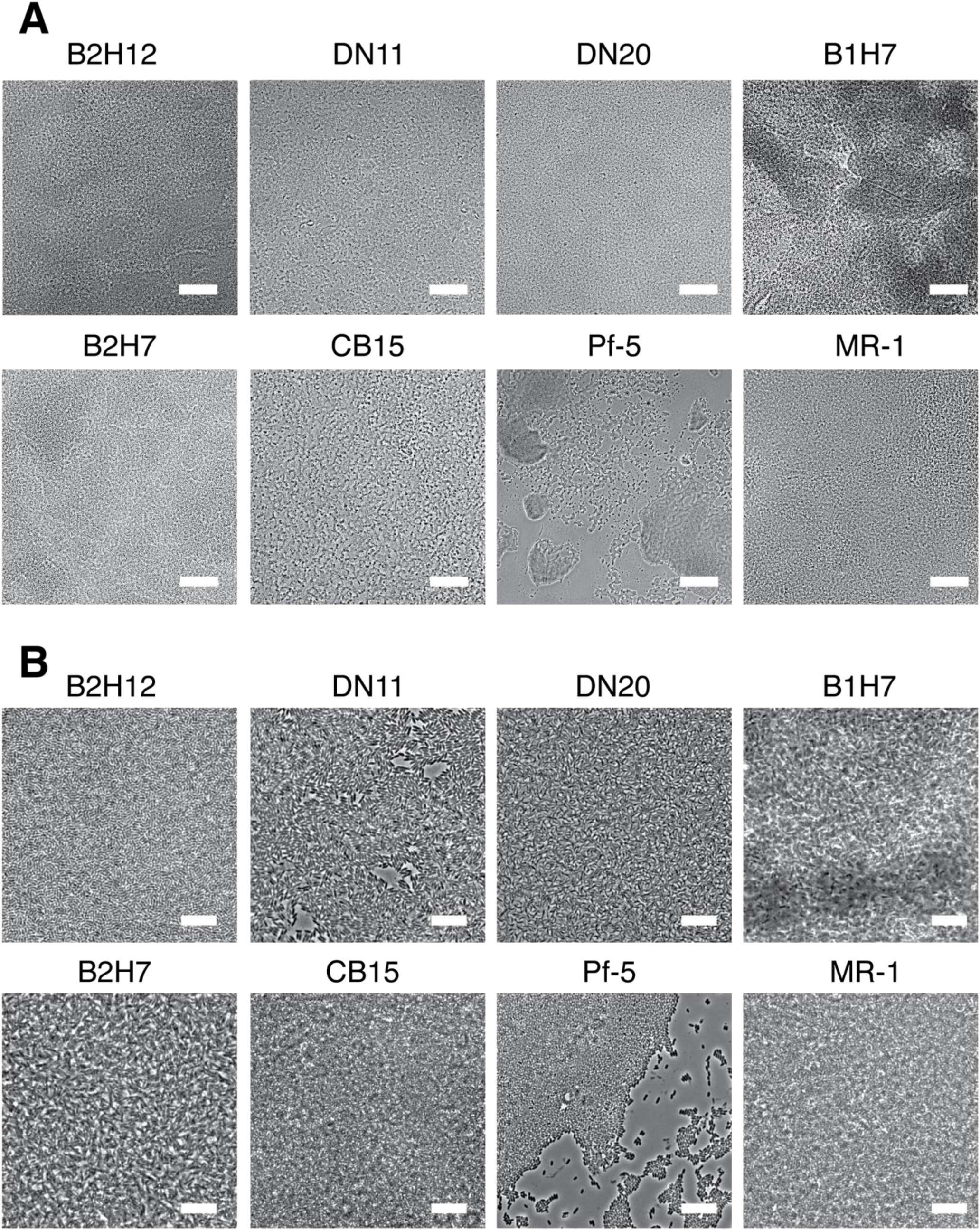
Phase contrast micrographs of extracted ALIs at 10X (A, scale bar represents 50µm) and 100X (B, scale bar represents 2.5µm) magnification. The ALIs from all cultures contains cells in complex, multi-layered structures. Note that the continuous Pf-5 pellicle layers were easily disrupted during extraction due to their inherent stiffness.

### Analysis of biofilm matrices from Pararheinheimera isolates

We extracted the extracellular matrices secreted by *Pararheinheimera* isolates in two ways. First, we centrifuged shaking liquid cultures to collect secreted products from the supernatant (Figure 5A). B2H12 and B1H7 cultures sedimented with a distinct pattern. A gelatinous layer was observed above the cell pellet, suggesting that these two strains secreted an insoluble matrix into the medium during growth. The remaining strains sedimented as a single cell pellet below spent liquid medium. Supernatants were collected and mixed with SDS to denature proteins and dissolve the gelatinous layer in the supernatants from B2H12 and B1H7. In parallel, we extracted ALI-associated biomass using steel mesh (Figure 3C) and dissolved the filtrate in SDS.

**Figure 5:**
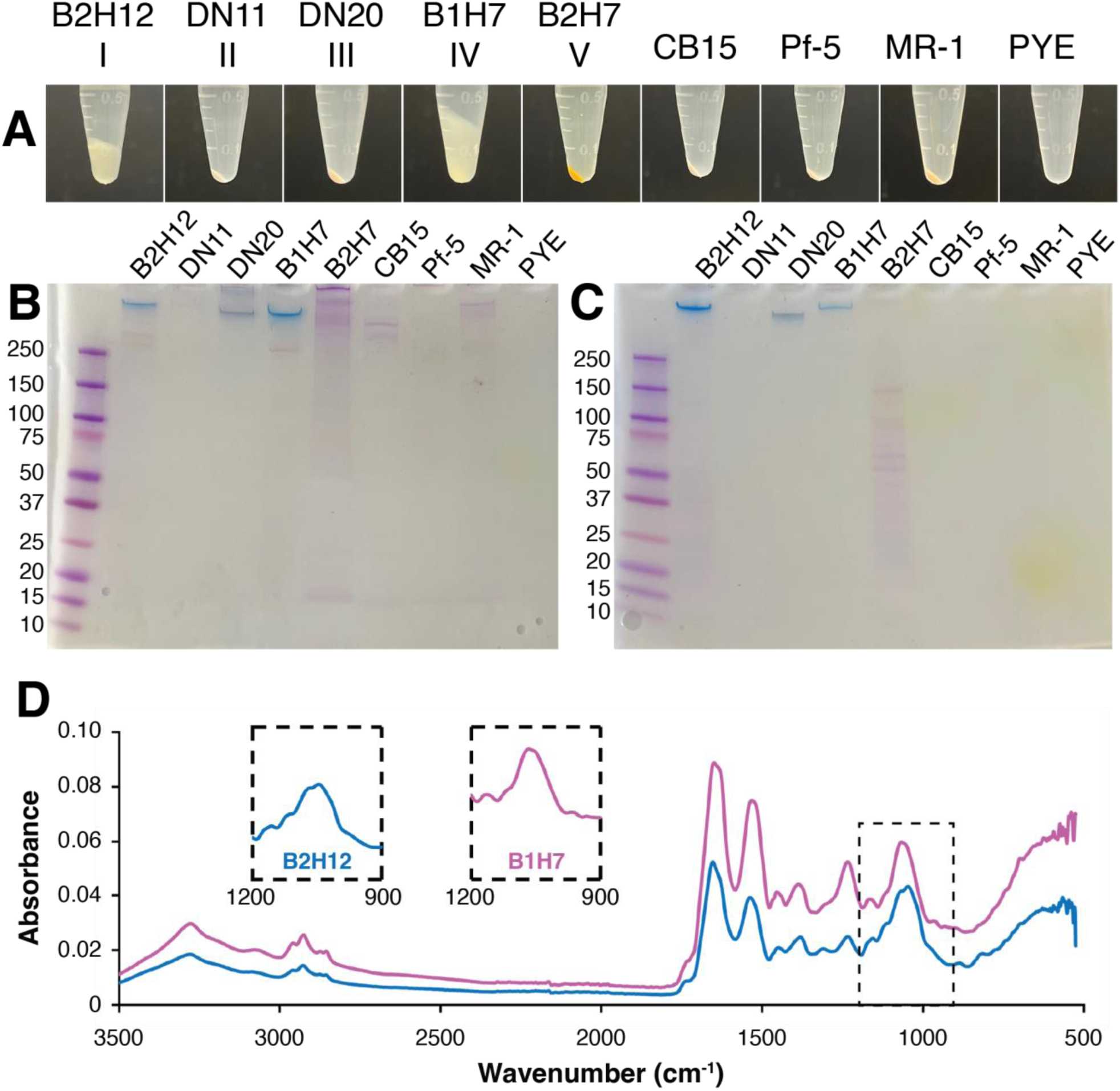
Biochemical analysis of secreted matrices from ALI colonizing isolates. (A) Centrifuged liquid cultures showing the production of an insoluble matrix in *Pararheinheimera sp.* B2H12 and B1H7. (B) SDS-PAGE analysis of steel mesh filtrates from static cultures of ALI colonizing bacteria. (C) SDS-PAGE analysis of supernatants from shaking cultures of ALI colonizing bacteria. (D) FTIR spectra of secreted matrices from B2H12 (blue) and B1H7 (magneta). A detailed view of the polysaccharide fingerprint region (1200-900cm^-1^) of each spectrum is shown in the insets.

The steel mesh filtrates (Figure 5B) and culture supernatants (Figure 5C) were analyzed by SDS-PAGE. Gels were stained with Stains-all, a cationic carbocyanine dye that stains proteins as dark bands that range from violet or deep blue to red and stains acidic polysaccharides as bright blue bands(43, 44). The supernatants from B2H12 (clade I) and B1H7 (clade IV) each displayed a high molecular weight (HMW, >250kDa) bright blue band indicative of acidic polysaccharides. The DN20 (clade III) supernatant contained a deep blue HMW band, and the supernatant from B2H7 (clade V) contained violet bands spanning a range of molecular weights. No signal was observed in the remaining supernatant extracts. The staining patterns of steel mesh extracts resembled the supernatant samples but with an increased signal for protein. The DN20 filtrate contained only the HMW blue band seen in the supernatant sample, and B2H7 filtrate contained violet protein staining that was more pronounced compared to the supernatant sample. B2H12 and B1H7 filtrates contained an additional violet HMW signal accompanying the bright blue bands seen in the supernatant samples. Filtrates from CB15 and MR-1 contained prominent violet colored protein signals that were not present in the respective supernatant samples. DN11 and Pf-5 filtrates contained faint violet staining suggestive of protein.

We were particularly interested in the insoluble matrices secreted by the viscous mass (VM) forming strains B2H12 and B1H7. SDS-PAGE analysis suggested that the gelatinous masses that accumulate at the ALI in these cultures contained an acidic exopolysaccharide (EPS) and protein. We extracted high-molecular weight compounds from the supernatants of B2H12 and B1H7 cultures (Materials and Methods) and analyzed the matrices using Fourier Transform Infrared Spectroscopy (FTIR, Figure 5D). Spectra for both matrices dissolved in water showed strong Amide I (1647 cm^-1^) and II (1531 cm^-1^) bands that are indicative of the presence of protein. Numerous peaks were also observed in the fingerprint region (1200-900 cm^-1^) for polysaccharides (Fig 5D). These results support the inference that the matrices secreted by B2H12 and B1H7 contain a mixture of exopolysaccharides (EPS) and proteins.

### Viscous mass (VM) biofilms contain thermally stable hydrogels

We analyzed the thermomechanical properties of the matrices secreted by B2H12 (clade I) and B1H7 (clade IV) to gain additional insight into the gelatinous biofilms produced by these strains. Thermogravimetric analysis (TGA) showed that both extracted matrices displayed high thermal stability. Both matrices had similar decay curves (Figure 6A). A minor loss of weight (∼7%) attributed to the loss of bound water was observed below 100°C. The major loss occurred in a range from 225-500°C for B2H12 and 200-500°C for B1H7, with midpoint values of 300°C. 20-25% residual mass remained stable past 600°C. Matrix extracts dissolved in water (1% w/v) were also examined using parallel plate rheometry. Both solutions displayed shear-thinning behavior, with the B2H12 matrix displaying higher viscosity than B1H7 at all shear rates (Figure 6B). The extracts also behaved as hydrogels at low strains and a frequency of 1 rad/s (Figure 6C). Linear viscoelastic behavior was observed at strains of up to 1.6% for B1H7 and up to 6.6% for B2H12. Frequency sweeps conducted at 1% strain confirmed that both secreted matrices formed hydrogels that were stable up to shear frequencies of ∼10rad/s. Dynamic moduli calculated from frequency sweep measurements demonstrated that gels formed by B2H12 extracts were considerably (∼10 times) stiffer than those formed by B1H7 (Figure 6D). Our results indicate that the EPS-rich matrices secreted by *Pararheinheimera* isolates from clades I and IV are thermostable materials that form hydrogels in aqueous solution. The hydrogels formed by B2H12 were particularly notable for their stiffness, with storage moduli approaching 100Pa.

**Figure 6:**
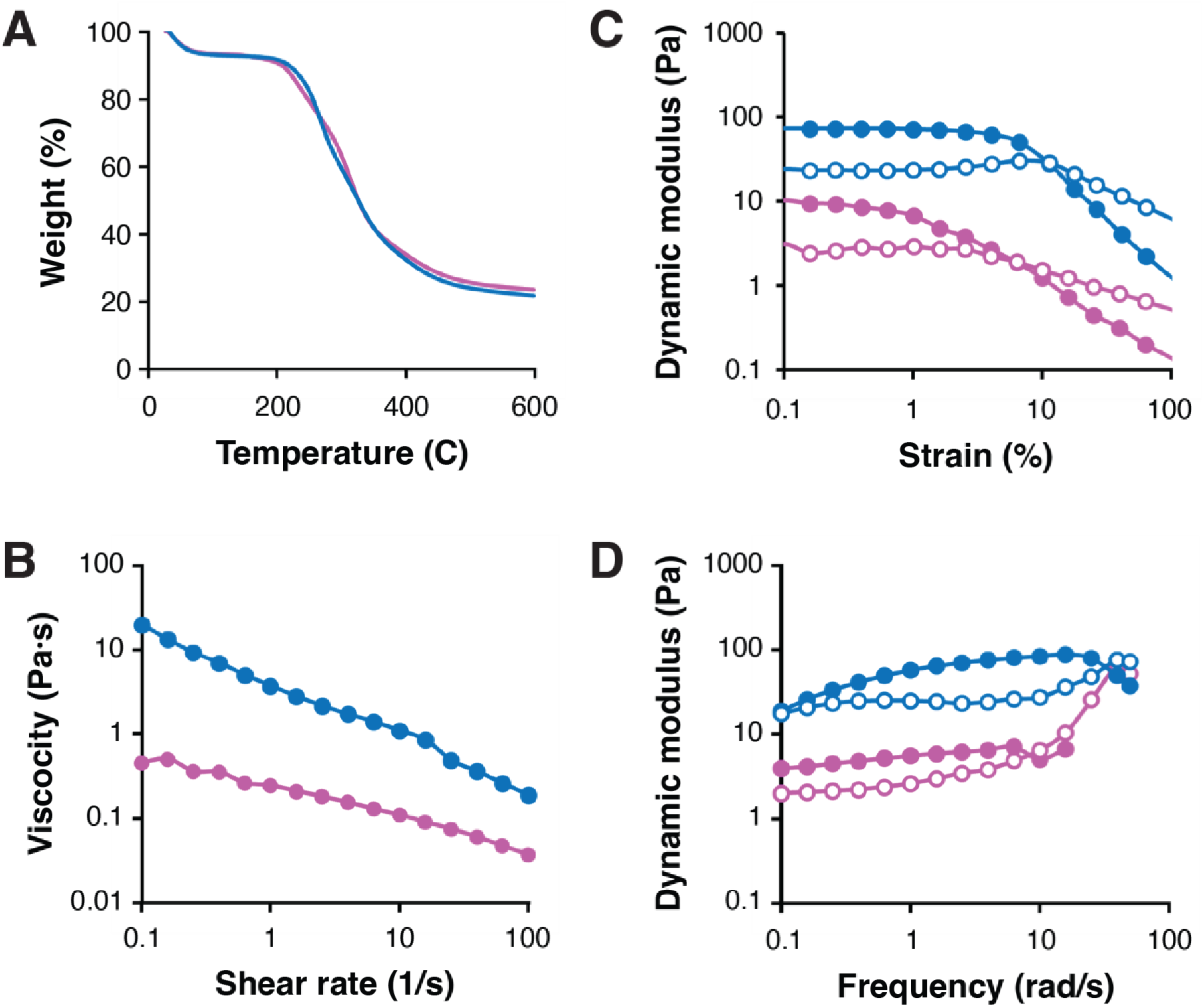
Thermomechanical properties of extracellular matrices from VM pellicle forming *Pararheinheimera* isolates B2H12 (blue) and B1H7 (magenta). (A) Thermogravitometric analysis of secreted matrices from strains B2H12 and B1H7. (B) Shear-thinning behavior of B2H12 and B1H7 matrices dissolved in water. Strain (C) and frequency (D) sweeps show that B2H12 and B1H7 matrices form stable hydrogels in solution. 1% (w/v) solutions were prepared in water for all rheology experiments.

## Discussion

The air–liquid interface (ALI) is a universal feature of aquatic ecosystems, but the strategies microbes use to thrive in this niche remain poorly characterized. The goal of this study was to explore how aquatic microbes persist at the ALI. We showed that microbes from freshwater can rapidly colonize the air–liquid interface when provided with conditions that support growth. The consistent emergence of stable pellicles during our enrichment experiments highlights the selective advantages associated with ALI occupancy. Although only a subset of lake-associated microbes proliferated under these conditions, the resulting pellicles were multispecies and ecologically dynamic. The succession of early (*Pararheinheimera*, *Shewanella, Aeromonas*), middle (*Janthinobacterium*) and late (*Stenotrophomonas*) colonizers, reinforces how ALI biofilms encompass the combined activities of multiple organisms. Members of the pellicle community employ distinct colonization strategies and produce biofilm matrices with different physical properties. Contributions from multiple taxa are likely to increase the complexity of the biofilm matrix, and future studies that explore the spatial and mechanical properties of these communities could illuminate how community dynamics influence ALI colonization.

The *(Para)rheinheimera* clade is distributed across diverse environments, including freshwater, marine, soil, host-associated and engineered habitats(45–50). However, little is known about the physiology or biofilm formation behavior of these organisms. *Pararheinheimera* strains accumulated in a reproducible pattern during pellicle enrichment. Although present at detectable levels in lake water, *Pararheinheimera* rapidly accumulated at the air–liquid interface during mixed-species pellicle growth before declining to undetectable levels. The transient accumulation at early stages of pellicle maturation was further supported by isolation experiments. *Pararheinheimera* strains were isolated directly from lake water and from pellicles harvested during the first four days after inoculation but were never recovered at later stages (Figure 1 and Table 1). Isolates grew rapidly and efficiently colonized the ALI in monoculture, demonstrating their intrinsic capacity for rapid surface access (Figure 3). We speculate that *Pararheinheimera* strains may function as early colonizers that help establish biofilms at the ALI. More broadly, our results point to a role for these organisms in structuring aquatic biofilms and motivate further investigation into the biofilms by this group.

While isolates from all five *Pararheinheimera* subclades colonized the ALI efficiently, they showed substantial variation in biofilm architecture. B2H7 (clade V) formed a thin, adherent pellicle resembling the PC pellicles produced by model organisms, but the remainder of the isolates we examined were either non-adherent (clades I-III) or weakly adherent (clade IV). The ALIs from clade II and clade III cultures contained a thin layer of cells resembling PC structures but lacking the characteristic adhesive properties. In contrast, the non-adherent biofilms from clades I and IV presented as gelatinous masses that were cohesive enough to be filtered from liquid using steel mesh. The architectures we observed resemble the physically-cohesive (PC, clade V), floccular material (FM, clades II and III) and viscous mass (VM, clades I and IV) pellicles described in *Pseudomonas fluorescens* isolates by Speirs and coworkers(51). The VM phenotype was polyphyletic within the *Pararheinheimera* genus (Figure 2), and the broader variation seen among our isolates indicates that biofilm architectures can diversify rapidly among closely related strains.

The VM pellicle forming *Pararheinheimera* isolates B2H12 (clade I) and B1H7 (clade IV) both displayed mucoid colony morphologies on solid media and secreted insoluble material during growth in shaking liquid cultures (Figures 2 and 5). The correlation among these phenotypes suggests that robust matrix secretion is an intrinsic feature of VM pellicle formation. FTIR analysis and differential staining of extracts from these strains demonstrate that both matrices contain a substantial fraction of acidic exopolysaccharide along with associated proteins. When these matrices were reconstituted in water, they formed thermally stable, shear-thinning hydrogels (Figure 6). However, matrix from the clade I isolate B2H12 formed gels that were approximately an order of magnitude stiffer (G’ of ∼70Pa, Fig6C) than that from the clade IV isolate B1H7 (G’ of 9Pa, Figure 6C). Detailed compositional and structural analysis of these matrices will be needed to understand the molecular basis for their distinct material behavior.

The VM biofilms produced by *Pararheinheimera* isolates show similarities to mucoid variants of *Pseudomonas aeruginosa*. Adaptive mutations that cause overproduction of the acidic polysaccharide alginate induce mucoidy in *P. aeruginosa*(52). Although few studies have directly examined their ALI colonizing behavior, viscoelastic moduli measured for surface-grown mucoid biofilms with varying alginate acetylation levels span the same range (1-100Pa) observed for B2H12 and B1H7 extracts(53). Bulk rheology with surface-grown *P. aeruginosa* indicates that mucoid biofilms are less stiff than non-mucoid forms, potentially due to the reduced cell densities resulting from increased matrix production(54). However, direct comparisons between VM- and PC-type pellicles are methodologically challenging. Interfacial rheology, uniaxial loading and other techniques applied to probe thin, PC-type pellicles would be poorly suited for the bulky, non-adhesive VM-type pellicles described here(27, 36, 55). VM and PC pellicles seem to represent fundamentally different materials.

The cellular mechanisms underlying PC-type pellicle formation have been established. Flagellar motility and chemotaxis help guide cells toward the oxygen-rich surface layer but are not strictly required to form PC pellicles(39, 56, 31). Production of the biofilm matrix is the crucial determinant of ALI colonization(29, 27, 31). Surface tension and other interfacial forces impart solid-like properties to the ALI that seem to facilitate direct attachment to the interface(57, 58). The aquatic bacterium *Caulobacter crescentus* can adhere directly to the surface boundary using a dedicated adhesin(31). Other studies propose that matrix driven adhesion to the walls of culture vessels provides a nucleation point for subsequent spread across the ALI(59). These models imply that the upper layer of the water column serves as a solid-like substrate for structures that resemble traditional biofilms. However, direct adhesion to the interface seems unlikely to explain the formation of VM biofilms. VM pellicles are inherently non-adherent, and they appear suspended near the surface rather than in direct contact with the ALI (Figure 3). We propose that copious secretion of gelatinous EPSs generates hydrated matrices with sufficient buoyancy to maintain position near the surface. Similar principles drive the ecology of flocs, transparent exopolymer particles (TEPs), marine snow and other polymer-rich aggregates in aquatic systems(60). Further investigation of VM-forming *Pararheinheimera* may provide insight into the role of buoyant matrices in shaping the dynamics of air–water interfaces.

## Materials and Methods

### Growth of mixed-species biofilms from freshwater samples

Water samples were collected from the south shore of Lake Mendota in Madison, WI during the summer and autumn months (May–October) of 2021-2025. A 1L graduated cylinder was inserted into the water to collect ∼500mL of surface water. Large particles were allowed to settle in the graduated cylinder for 1hr before either directly plating water on agar plates to isolate bacteria or inoculating into 20 x 150mm glass culture tubes to grow mixed species pellicle biofilms. Peptone-yeast extract (PYE) medium (0.2% w/v peptone, 0.1% w/v yeast extract, 1mM MgSO_4_, 0.5mM CaCl_2_) was used as a nutrient source for all cultivation experiments. Four different growth conditions were used to enrich pellicles: (1) 20mL of lake water sample; (2) 2mL of lake water, 17.9mL sterile H_2_0, 0.1mL PYE medium (3) 2mL of undiluted lake water, 17mL sterile H_2_0, 1mL PYE medium (4) 2mL of undiluted lake water, 18mL PYE medium. Cultures were allowed to incubate statically at room temperature under ambient light for up to one month with periodic sampling from the ALI.

### Microscopy

ALI samples were extracted using a modified version of the method described by Aretha Fiebig(31). Briefly, the large end of a P1000 pipette was used to excise a circular section from the ALI. The pipette was submerged 1-2 mm and when removed was covered in the film, leaving behind a scar in the pellicle. A similar technique was used for the *Pararheinheimera* isolate pellicle samples, with the use of the large end of a glass 5 mL serological pipette for excising a circular plug instead of a P1000. The film was transferred to a thin agar pad on a glass slide by gently pressing the large end of the pipette to the pad and lifting so as not to wrinkle the sample. The film was left to dry on the slide for five minutes and then a coverslip was placed on the sample. Microscopy was performed using a Nikon Ti-E inverted microscope, and images were captured with an Orca Fusion BT digital CMOS camera (Hamamatsu).

### 16S amplicon profiling of mixed-species biofilms

Sampling of the ALI for DNA isolation was performed as described above. In place of transferring the film to a glass slide, the large end of the pipette (P1000) was transferred to a 5 mL centrifuge tube with 3 mL of PYE. The tube was centrifuged at 6000 RCF for 5 minutes to collect only the ALI film in the pellet. This pellet was resuspended in 800 µL of CD1 or lysis buffer from the Qiagen PowerSoil Pro Kit and transferred to PowerBead Pro tubes. Samples were heated to 65°C for 5 minutes before being vortexed at max speed for 5 minutes. To sample the lake water pre-enrichment, 200mL of lake water was filtered through a 0.2µm Pall Supor PES membrane disc filter. The filter was added to a PowerSoil bead beating tube with 800 µL of CD1 buffer and was vortexed at max speed for 5 minutes. The tube was centrifuged at 4000xg for 1 minute and the contents of the buffer were collected while leaving behind the filter. All samples were further processed as described in the Qiagen PowerSoil Pro Kit.

Amplicon libraries were prepared and analyzed as described in Arellano et al. (61). Briefly, the V4 hypervariable region of the 16S rRNA gene was amplified using a one-step amplification with dual-indexed universal bacterial primers 515F (5′-GTGCCAGCMGCCGCGGTAA-3) and 806R (5′-GGACTACHVGGGTWTCTAAT-3)(62). Prior to amplification samples were diluted to ∼10 ng/µL. Amplification was confirmed for all samples using gel electrophoresis (1% agarose), prior to library purification using a MagJET NGS Cleanup and Size Selection Kit (Thermo Scientific). Resulting libraries were quantified and pooled at equimolar concentrations followed by paired-end sequencing (2×250-bp) on an Illumina MiSeq instrument.

On-board sequence demultiplexing was conducted on the Illumina MiSeq instrument using the Generate FASTQ Analysis Module. Subsequent quality control was conducted using the QIIME2 command-line platform (2024.5)(63). The tool DADA2 was used to denoise parallel reads before read-joining(64). Forward reads were not truncated due to a high q-score throughout, but reverse reads were truncated to 140 base pairs. The QIIME2 Greengenes2 (2022.10) package was used to assign taxonomy to reads(65). A phylogenetic tree for the reads was prepared and re-rooted using the code described by Joey711 on github. A decontamination step was run using the R package “decontam”, comparing samples to PCR and extraction controls(66). In this step, chloroplasts and mitochondria were also removed from the 16S dataset. The R package “ggplot2” and its extension “ggh4x” were used to make abundance plots.

### Isolation of Pararheinheimera strains

Samples collected from the ALI of enrichment cultures were resuspended in PYE medium and homogenized thoroughly by pipetting. Each was serially diluted in PYE and spread on PYE agar plates. Plates were incubated at room temperature for one week. Colonies were scraped from plates based on their morphologies re-streaked three times successively on PYE agar. Isolates were then grown in PYE medium with 200rpm shaking at 25°C before adding glycerol and storing at −80°C.

### Phylogenetic analysis

Cells scraped from agar plates were suspended in 20µL of ThermoPol buffer (20 mM Tris-HCl pH 8.8, 10 mM (NH_4_)_2_SO_4_, 10 mM KCl, 2 mM MgSO_4_, 0.1% Triton X-100) and heated at 95°C for 10m. 2µL of the resulting lysate was used as a template for PCR amplification of the 16S rDNA using the primers 27F and 1492R. PCR products were treated with ExoSap-IT and sequenced using the Sanger method. Assembled 16 rDNA sequences were aligned using CLUSTALW and phylogenetic trees were inferred using the Maximum Liklihood method within the MEGA suite(67).

### Analysis of static biofilm formation

One representative from each of the five *Pararheinheimera* clades was chosen and evaluated for pellicle formation. Overnight cultures of each isolate were diluted to 0.05 OD_660_ in 4 mL of PYE. Cultures were left to incubate statically for 3 days at room temperature in a humidity chamber. For the humidity chamber, two beakers were filled with 35g of KNO_4_ and 100 mL of Milli-Q water. These beakers were placed in a large, lidded plastic bin with the samples. After 3 days, pellicles were observed and photographed. Tubes were then decanted, 1 mL of Milli-Q water was added to each tube, and tubes were vortexed at medium setting for 10 seconds. Liquid was decanted and 4 mL of 0.01% crystal violet was added to the tubes and left to incubate statically for 10 minutes. The dye was discarded, and 1 mL of Milli-Q water was added to all tubes and vortexed again for 10 seconds at medium setting, then decanted. Tubes were photographed. Other bacteria that are known for forming pellicles were used as controls to compare biofilm formation. For this, *Caulobacter crescentus* CB15, *Pseudomonas protegens* Pf-5, and *Shewanella oneidensis* MR-1 were used and grown/treated as described for *Pararheinheimera* isolates.

### Biofilm matrix extraction

A representative from each of the five *Pararheinheimera* clades as well as pellicle control organisms mentioned above were evaluated for matrix characteristics. Overnight cultures of all five *Pararheinheimera* isolates as well as the three controls were diluted to 0.05 OD_660_ in 3.5 mL of PYE in a 12-well plate. Stainless steel mesh was prepared and placed at the bottom of the wells in a 12-well plate as described in Pisithkul et al. 2019(68). Briefly, steel mesh was cut to fit inside of the wells of a 12-well plate with overhangs that rested on the tops of the well, allowing floating biofilms to be extracted from the wells by lifting the steel mesh. Plates were incubated statically for 3 days in a humidity chamber before the steel mesh was lifted and photographed. For SDS-PAGE analysis of steel mesh samples, steel mesh was lifted and placed into a new 12-well plate with 500 µL of 2% SDS and incubated shaking at room temperature for 3 hours.

Large EPS extractions were carried out for rheometric and thermogravimetric analyses. For this, 400 mL of PYE were inoculated with 3 mL of overnight cultures of *Pararheinheimera* B2H12 and B1H7. The B2H12 culture incubated for three days while the B1H7 culture incubated for two days, both shaking at room temperature. Both 400 mL cultures were centrifuged in a Sorvall RC 5B centrifuge at 4200 RPM for 45 minutes at 4°C. The media was decanted and the gelatinous EPS pellets were carefully pipetted away from the dense cell pellet. EPS pellets were transferred to a 50 mL conical tube and centrifuged again, leaving behind an additional small cell pellet, in a swinging bucket Sorvall X Pro Series centrifuge at 4815 x g for 20 minutes at room temperature. The liquid was decanted, and 8 M urea was added at the same volume as the remaining EPS pellets (1:1). Samples were mixed until the EPS was dissolved and then transferred to Spectra/Por7 pre-treated 50 kD MWCO dialysis tubing and dialyzed against Milli-Q water overnight. The samples were moved to fresh water for three more iterations of dialysis against Milli-Q water and then aliquoted into glass tubes and lyophilized.

### SDS-PAGE analysis of biofilm matrices

For preparing samples, 2x Laemmli buffer was pre-warmed and mixed with samples at a 1:1 dilution. Samples were then placed on a 65°C heating block for 15 minutes before 12.5µL were loaded on to gels. All PAGEs were run using mini-PROTEAN TGX pre-cast gels with a gradient of 4-20% (cat. # 4561093). The running buffer used was a 1x tris glycine buffer, and gels were run for an hour at 118 V.

PAGEs were stained using stains-all. Gels were pre-treated twice in 25% (v/v) isopropanol for 1.5 hours. Stains-all stain was prepared in glass vessels protected from light(69). 5 mL of Stains-all dissolved at 0.1% (w/v) in dimethyl formamide (DMF) was added to 5 mL of DMF, 25 mL of isopropanol, 5 mL of 300mM tris HCl pH 8.8, and 60 mL of water to prepare the stain. PAGEs were stained overnight on a rocking incubator and destained the following morning by moving the PAGE to water and incubating for about an hour in light(70).

### Thermogravimetric analysis

Thermogravimetric analysis was completed on a TA Instruments 5500 thermogravimetric analyzer (Texas Instruments, Dallas, TA) following the protocol developed for the promonan EPS from *Sphingomonas sp.* LM7(71, 72). Samples were initially weighed and loaded using a platinum pan. Weights were adjusted to between 2-10 mg. The ramp rate was set to 10 °C/min and the gas supply was set to nitrogen.

### Rheometry

1% (w/v) solution of each extracellular matrix were prepared by dissolving lyophilized extracts in deionized water. Experiments were conducted using a TA Instruments DHR3 rheometer (Texas Instruments, Dallas, TA). A strain sweep at a frequency of 1 rad/s was used to determine the linear viscoelastic region and select a strain of 1% to be used for frequency sweeps. Both a strain sweep and a frequency sweep were conducted for each sample. A 20 mm parallel plate fixture was used with the gap set to 300 μm, requiring a sample volume of approximately 100 μL. The fixtures were thoroughly cleaned before use and between samples with deionized water and isopropyl alcohol and dried with compressed air. The sample was pipetted onto the bottom plate and the upper fixture lowered to the set gap value. A visual check confirmed that the amount of sample adequately filled the gap. Graphs presented are the average of three independent trials. Sweeps of the strain rate were used to produce the viscosity and stress graphs (n=1).

## Data availability

16S amplicon sequencing datasets have been submitted to the NCBI Sequence Read Archive (SRA) under BioProject ID PRJNA1400529. 16S rDNA sequences for all 31 *Pararheinheimera* isolates were deposited in the NCBI GenBank as accessions PX310746-PX310776 (Table 1).

## Acknowledgements

This work was supported by a Beckman Young Investigator award to D.M.H., National Institutes of Health award R35GM150652 to D.M.H, the NSERC PGS-D scholarship (E. W. v. W.). It was also supported in part and performed in part at the Engineered Living Materials Institute at Cornell University and the Cornell University Materials Research Science and Engineering Center.

